# The DEAD-box protein p68 and β-catenin: the crucial regulators of FOXM1 gene expression in arbitrating colorectal cancer

**DOI:** 10.1101/2022.10.28.514256

**Authors:** Shaheda Tabassum, Malini Basu, Mrinal K Ghosh

## Abstract

Forkhead box M1 (FOXM1), a vital member of the Forkhead box family of transcription factors, helps in mediating oncogenesis. However, limited knowledge exists regarding the mechanistic insights into the FOXM1 gene regulation. p68, an archetypal member of the DEAD-box family of RNA helicases, shows multifaceted action in cancer progression by arbitrating RNA metabolism and transcriptionally coactivating transcription factors. Here, we report a novel mechanism of alliance between p68 and the Wnt/β-catenin pathway in regulating FOXM1 gene expression and driving colon carcinogenesis. Initial bioinformatic analyses highlighted elevated expression levels of FOXM1 and p68 in colorectal cancer datasets. Immunohistochemical assays confirmed that FOXM1 showed a positive correlation with p68 and β-catenin in both normal and colon carcinoma patient samples. Overexpression of p68 and β-catenin increased the protein and mRNA expression profiles of FOXM1, and the converse correlation occurred during downregulation. Mechanistically, overexpression and knockdown of p68 and β-catenin elevated and diminished FOXM1 promoter activity respectively. Additionally, Chromatin immunoprecipitation assay demonstrated the occupancy of p68 and β-catenin at the TCF4/LEF binding element (TBE) sites on the FOXM1 promoter. Thiostrepton delineated the effect of FOXM1 inhibition on cell proliferation and migration. Colony formation assay, migration assay, and cell cycle data reveal the importance of the p68/β-catenin/FOXM1 axis in oncogenesis. Collectively, our study mechanistically highlights the regulation of FOXM1 gene expression by p68 and β-catenin in colorectal cancer.

## Introduction

DEAD-box RNA helicase p68 (DDX5) is an ATP-dependent RNA helicase that is intricately linked with cancer, owing to its helicase activity in RNA metabolism and coactivating numerous genes such as Androgen receptor (AR), β-catenin, p53, Runx2, MyoD, etc. It is overexpressed in most cancers, colon cancer being the most implicated. It contributes to colon cancer through up-regulation of proto-oncogenes and down-regulation of tumor suppressors. Thus, it is a pivotal member involved in carcinogenesis and needs to be comprehensively studied to provide insights into the process of oncogenesis and tackle the menace of tumor progression^1–4^.

The Wnt signaling pathway plays an integral role in regulating cell cycle, developmental pathways, and inflammatory signaling axis. The classical canonical Wnt pathway helps in differentiation, proliferation, and survival whereas cell polarity and migration are regulated by a non-canonical pathway. The canonical Wnt/β-catenin signaling pathway is highly implicated in many cancers. Moreover, mutations of the Wnt/β-catenin pathway are often recognized as causative factors for most of the colorectal cancer (CRC) cases. β-catenin, an effector component of this pathway, is gaining prominence as a molecular marker of cancer. Therefore, understanding the functionality of this pathway is indispensable for developing therapies for colon cancer^5,6^.

In terms of morbidity and mortality, CRC is the leading malignant tumor globally. It is ranked third in terms of worldwide incidence and second in terms of mortality rates among malignant tumors. Current therapies include endoscopic and surgical local excision, radiotherapy, systemic therapy, extensive surgery for metastatic disease, local ablative therapies, palliative chemotherapy, targeted therapy, and immunotherapy^7,8^. However, the challenges of metastasis pose considerable problems in the therapeutic arena. Accumulation of genetic mutations, aberrant cellular signaling pathways, heightened expression of oncogenes, and suppression of tumor suppressor genes together contribute to the invasive carcinoma condition. Apt and efficient screening programs can significantly reduce the risk through early detection and immediate administration of treatment modalities^9–12^.

The Forkhead box (Fox) family of proteins are evolutionarily conserved transcription factors that are composed of a monomeric forkhead DNA-binding domain (winged-helix domain). FOXM1, a crucial member of this family is popular for its role as a master regulator of the cell cycle and for promoting oncogenesis. It is overexpressed in different cancers being linked with cell differentiation, proliferation, migration, invasion, tissue homeostasis, repair of DNA damage, angiogenesis, and drug resistance during tumorigenesis^13–19^. Clinically, FOXM1 acts as a prognostic marker in colorectal cancer due to its high expression levels and poor survival rates. However, a lot remains to be explored regarding the mechanisms of FOXM1 gene regulation. p68 acts as a coactivator of β-catenin^20^ in several cancer types whereas a cross-talk between the Wnt/β-catenin pathway and FoxM1 has been reported in glioma stem cells^21–24^. Therefore, we sought to investigate the role of p68 and Wnt/β-catenin in the regulation of the FOXM1 gene expression in CRC and delineate the mechanistic details of this synergism. Understanding the molecular mechanisms will pave way to strengthen future research in the therapeutic area.

## Results

### Expression profiles of p68 and FOXM1 using bioinformatics analyses

Using UALCAN^25^, we generated a pan-cancer expression profile of FOXM1. It used mRNA expression data set in 24 different types of cancer and plotted them as log2 (TPM+1) vs TCGA samples and comparing normal with tumor samples. FOXM1 was found to be significantly overexpressed in different cancers (Figure 1a). A similar pan-cancer increased expression profile of p68 (DDX5) was obtained using UALCAN database (Figure 1b). Colorectal Adenocarcinoma (COAD) was chosen for further analysis. Using COAD dataset, the transcript levels of FOXM1 and DDX5 were checked in primary tumor vs normal samples, where a significant increase in the expression levels of FOXM1 and DDX5 in primary tumor samples was observed (Figs. 1c and 1d). Similarly, a concomitant increased expression profile of FOXM1 and DDX5 transcript levels with increasing grades of colorectal cancer progression was seen (Figs. 1e and 1f) thereby implicating their correlation bioinformatically.

**Fig 1.**
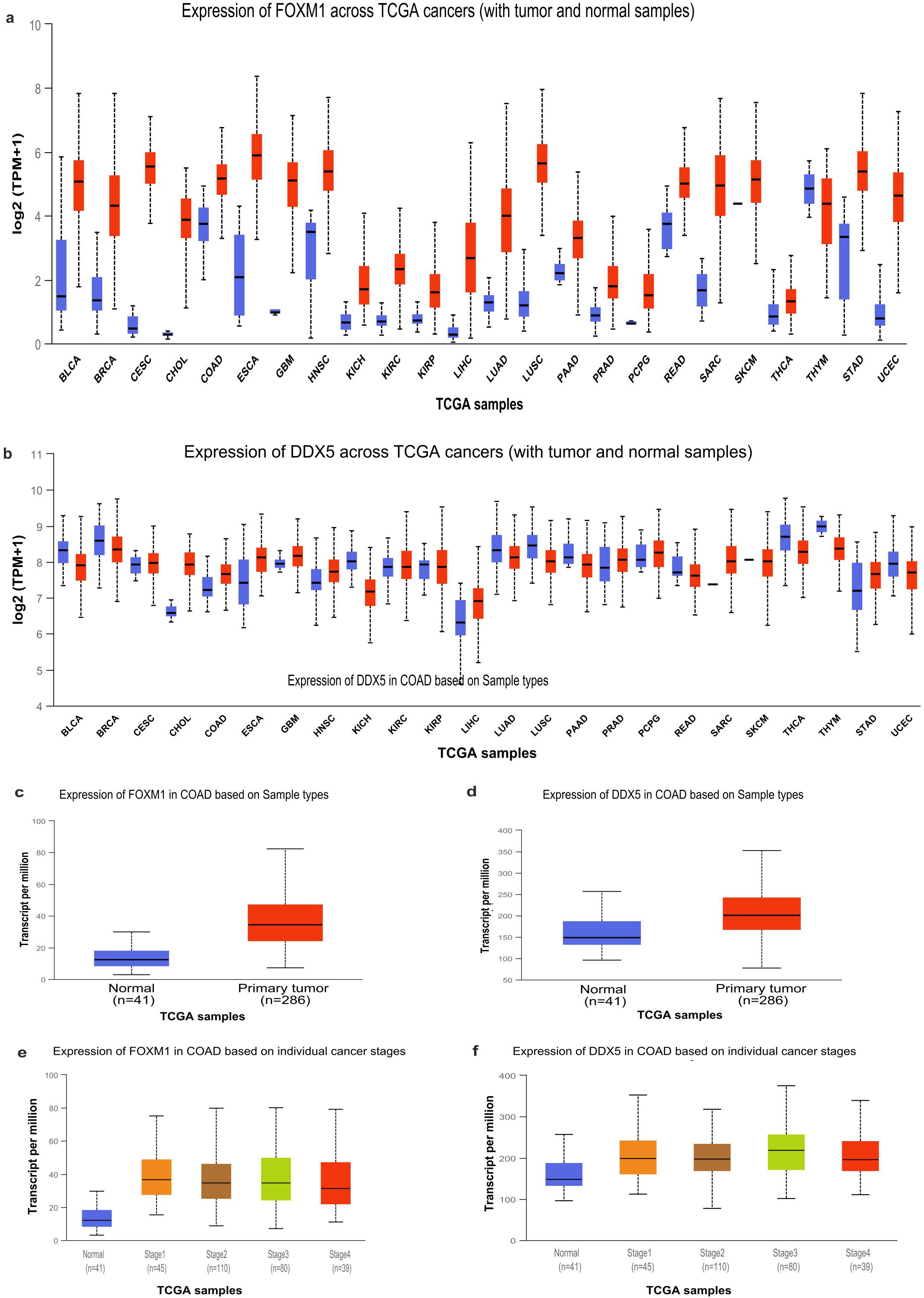
Bioinformatic analysis of expression profiles of FOXM1 and p68. **(a)** and **(b)**, UALCAN platform-based analysis of FOXM1 and p68 expression across different cancers (http://ualcan.path.uab.edu/analysis.html). BLCA, bladder urothelial carcinoma; BRCA, breast carcinoma; CESC, cervical squamous cell carcinoma, and endocervical adenocarcinoma; CHOL, cholangiocarcinoma; COAD, colon adenocarcinoma; ESCA, esophageal carcinoma; GBM, glioblastoma multiforme; HNSC, head and neck squamous cell carcinoma; KICH, Kidney chromophobe; KIRC, kidney renal clear cell carcinoma; KIRP, kidney renal papillary cell carcinoma; LIHC, liver hepatocellular carcinoma; LUAD, lung adenocarcinoma; LUSC, lung squamous cell carcinoma; PAAD, pancreatic adenocarcinoma; PRAD, prostate adenocarcinoma; PCPG, pheochromocytoma and paraganglioma; READ, rectum adenocarcinoma; SARC, sarcoma; SKCM, skin cutaneous melanoma; THCA, thyroid carcinoma; THYM, thymoma; STAD, stomach adenocarcinoma; **(c)** and **(d)**, The expression levels of FOXM1 and DDX5 was bioinformatically assessed in COAD tissue samples with that of normal tissue samples. **(e)** and (**f)**, representative study of FOXM1 and DDX5 expression levels through cancer stage-specific (I-IV) analysis of COAD tumor samples compared with normal tissue samples.

### FOXM1 and p68 are positively correlated in colon carcinoma tissue samples and p68 regulates FOXM1 expression in CRC cell lines

To explore the connection between p68 and FoxM1, immunohistochemical analyses (IHC) was performed in colon carcinoma (n = 20) and adjacent normal colon tissue samples (n = 6). p68 and FoxM1 exhibited elevated expression in colon carcinoma samples compared to normal colon samples (Fig. 2a). H-score quantification revealed significant staining intensity differences between p68 and FoxM1 and were represented using Notched Box Plot (Fig. 1b). Statistical significance was predicted by the Mann–Whitney U-test analysis. Furthermore, their mean H-score difference was found to be statistically significant in colon carcinoma samples compared to the normal samples (Fig. 1c). The intensity of staining shown by p68 and FoxM1, depicted by their respective H-scores, followed comparable trends in both normal and colon carcinoma samples (Fig. 1d). Finally, the Spearman’s rank correlation test confirmed that H-scores of p68 and FoxM1 bear a strong positive correlation (r_s_ = 0.9077) (Fig. 1e). Thus, the results hint a plausible connection and correlation between p68 and FoxM1.

**Fig 2.**
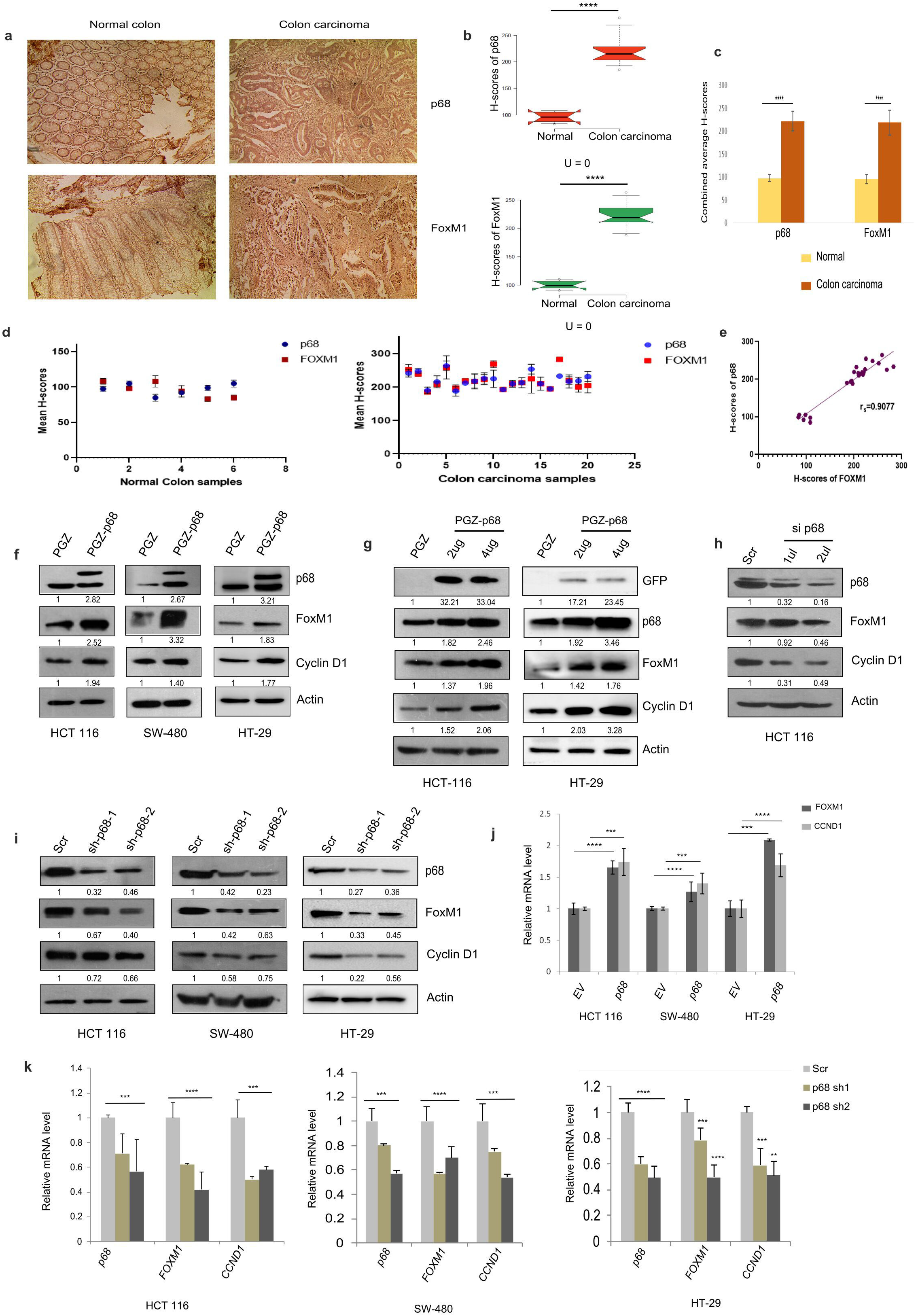
Positive correlation between FOXM1 and p68 in human colon carcinoma tissues and regulation of FOXM1 by p68 at protein and transcript levels. **(a)** Representative images from Immunohistochemical (IHC) staining of p68 and FOXM1 conducted in human normal colon and colon carcinoma tissue samples. The images were captured at 200X magnification. **(b)** The H-scores of p68 and FOXM1 were represented in the form of a Notched box plot in normal (n = 6) and colon carcinoma (n = 20) samples; H-scores were used for the calculation of Mann–Whitney U-values. **(c)** Comparative column graphs of the combined average H-scores of p68 and FOXM1. The mean (+/-) s.d is represented by error bars; the p-values were calculated using an independent, two-tailed Student’s t-test, and p < 0.0001 is signified as ****. **(d)** The mean H-scores of p68 and FOXM1 were compared with the help of scatter plots in normal and colon carcinoma tissue samples, respectively. **(e)** Spearman’s rank correlation coefficient (r_s_) was depicted graphically by using the mean H-scores of p68 and FOXM1 from both normal and colon carcinoma tissues. **(f)** Multiple colon cancer cell lines (HCT 116, SW-480, and HT29) were grown in 60 mm cell culture plates. They were transfected with 4 μg of pGZ-p68 or the empty PGZ vector and harvested after 24hrs. Immunoblot depicts the expression of p68, FoxM1, Cyclin D1, and Actin proteins. Densitometric analyses relative to Actin (loading control) are mentioned below the blots. **(g)** HCT 116 and HT29 cells, seeded in 60mm dishes, were transfected with sequentially increasing doses of pGZ-p68 against EV as indicated and thereafter respective proteins were immunoblotted. ‘Exo’ and ‘Endo’ portrays exogenous and endogenous p68, respectively. Densitometry values are mentioned below the images of the respective blots. **(h)** HCT 116 cells were transfected with successive doses (1 μl and 2 μl) of p68 siRNA along with control siRNA and harvested post 48hrs. Immunoblots of p68, FOXM1, Cyclin D1, and Actin are represented. 1 and 2 μl siRNAs were used from 10 μM stock **(i)** 4 μg of either scramble shRNA or p68 shRNA1 or p68 shRNA2 plasmids were transfected in multiple colon cancer cell lines and immunoblotting was performed after harvesting the cells post 48hrs from the experiment. Densitometry values are mentioned below the blots after comparison with a loading control. **(j)** cDNA prepared from PGZ-p68 or PGZ-transfected multiple colon cancer cell lines (HCT 116, SW 480, HT 29) were utilized in qRT-PCR analysis to compare and contrast the mRNA levels of genes of interest; FOXM1 and Cyclin D1 (CCND1) genes due to overexpression of p68. **(k)** In a converse approach, mRNA levels of p68, FOXM1, and Cyclin D1 genes were observed through qRT-PCR analysis after preparing cDNA from 35mm plates of multiple colon cancer cell lines transfected with 2 μg of either scramble shRNA or p68 shRNA1 and p68 shRNA2 plasmids and harvesting them post 48hrs from time of transfection. Cyclin D1 (CCND1) served as positive control whereas 18S rRNA was used for normalization. The mean (+/-) s.d is represented by error bars; the p-values were calculated using an independent, two-tailed Student’s t-test, and p < 0.0001 is signified as ****.

Furthermore, overexpression of p68 was done in multiple CRC cell lines (HCT 116, SW-480, and HT-29) that enhanced the protein levels of FoxM1. Cyclin D1 (the positive control) showed a similar trend (Fig. 2f). Next, p68 was overexpressed in a dose-dependent manner in HCT 116 and HT-29 cell lines that caused a concomitant dose-dependent increment of FoxM1 and Cyclin D1 protein levels (Fig. 2g). Conversely, endogenous levels of p68 were depleted using multiple small hairpin RNA (shRNA) and siRNA. Upon depleting p68 using siRNA in a dose-dependent manner, a sharp decline in the FoxM1 protein level was observed in the HCT 116 cell line with Cyclin D1 acting as a positive control^26^ (Fig. 2h). Similarly, upon downregulating p68 using shRNAs, a comparable decrease of FoxM1 was observed with Cyclin D1 following similar course (Fig. 2i). Further, to exemplify whether this connection exists at the transcript level, qRT-PCR analyses were conducted in multiple colon CRC lines (HCT 116, SW-480, and HT-29). Upon overexpressing p68, a concomitant increase in the transcript levels of the FOXM1 gene was observed (Fig. 2j). Conversely, upon downregulating p68 using multiple shRNAs, FOXM1 gene showed sharp declining transcript levels (Fig. 2k). Cyclin D1 served as the positive control while 18S gene served as an internal control. Also, decreasing trends of FOXM1 transcript level was observed upon downregulating p68 through siRNA-mediated knockdown in HCT 116 and HT-29 cell lines (Supplementary Figs. S1 a and b). Collectively, these results highlight a positive correlation between p68 and FOXM1 and emphasize that p68 regulates FOXM1 at both the protein and mRNA levels in colon cancer cell lines.

### Wnt/β-catenin signaling mediates FOXM1 gene expression in CRC

Wnt/β-catenin signaling is involved in colon carcinoma progression^6^. Moreover, p68 plays a major role in the coactivation of β-catenin to promote tumorigenesis ^27,28^. Also, FoxM1 promotes colorectal carcinogenesis and metastasis by activating the Wnt/β-catenin signaling^15,19,29^. Thus, we sought to explore the link between FoxM1 and β-catenin with p68 as a master regulator. Immunohistochemical analyses (IHC) were conducted and exhibited elevated expression of β-catenin and FoxM1 proteins in colon carcinoma samples (20) in contrast to normal samples (6) (Fig. 3a). Notched Box Plot was drawn to represent H-score quantification, which was statistically significant and differentiated the staining intensity of β-catenin between colon carcinoma and normal samples (Fig. 3b). The combined average H-score difference between β-catenin and FoxM1 proteins was found to be statistically significant in colon carcinoma samples (Fig. 3c). Comparable and correlated trends was obtained between the staining intensity of β-catenin and FoxM1 through their respective mean H-scores in normal and colon carcinoma samples (Fig. 3d). Furthermore, the Spearman’s rank correlation test confirmed a positive correlation between H-scores of p68 and FoxM1 (rs = 0.8353) (Fig. 3e). Therefore, β-catenin and FoxM1 proteins indicatively bear positive correlation in colon carcinoma samples.

**Fig 3.**
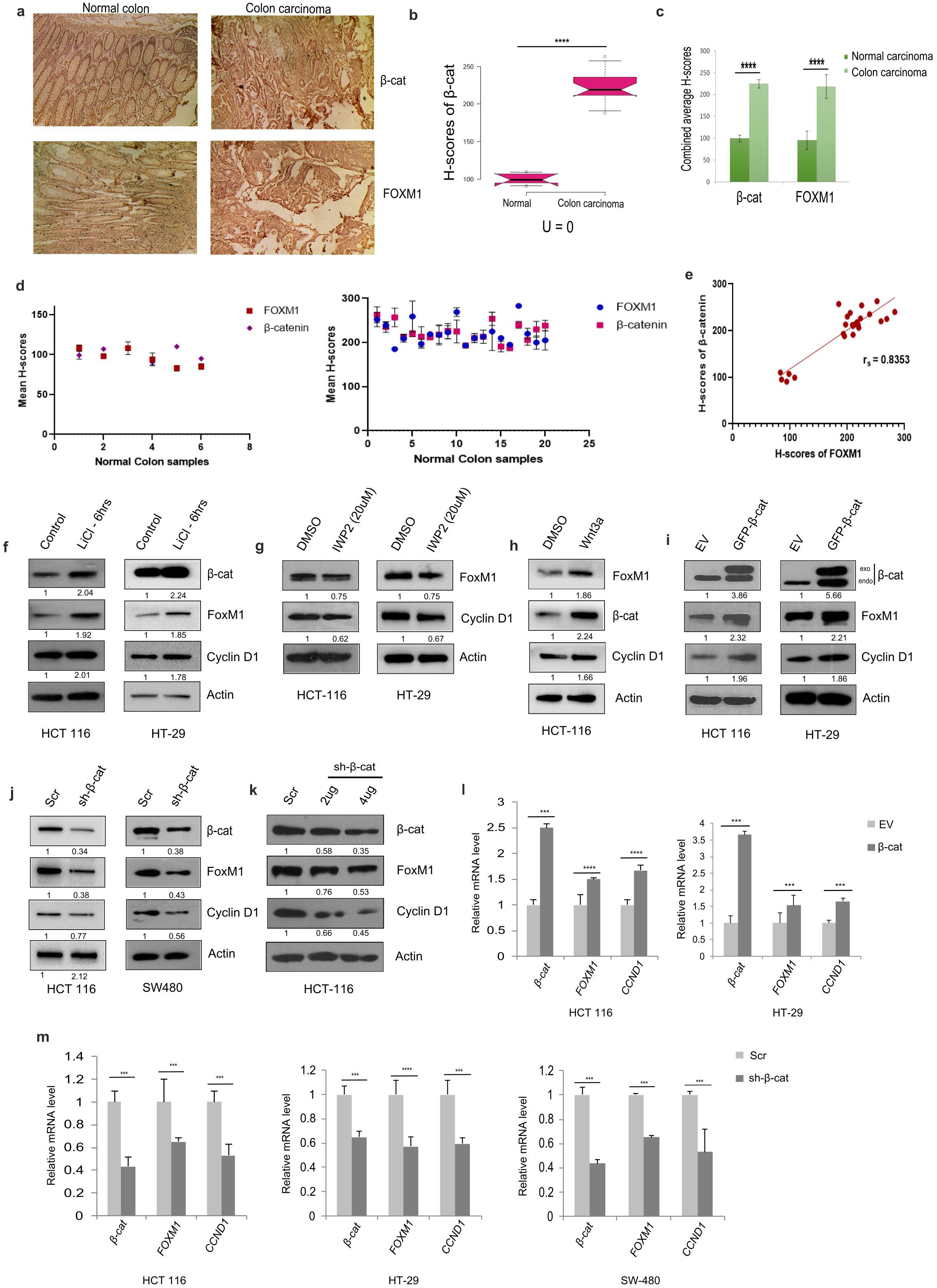
Involvement of Wnt signaling and mediation of regulation of FOXM1 by β-catenin. **(a)** Representative Immunohistochemical (IHC) staining images of β-catenin and FOXM1 were conducted in human normal colon and colon carcinoma tissue samples. The images were captured at 200X magnification. **(b)** Representative Notched-box plots were obtained from the H-scores of β-catenin and FOXM1 in normal (n = 6) and colon carcinoma (n = 20) samples; H-scores were further used for the calculation of Mann–Whitney U-values. **(c)** Comparison between the combined average H-scores of β-catenin and FOXM1. The mean (+/-) s.d is represented by error bars; the p-values were calculated using an independent, two-tailed Student’s t-test, and p < 0.0001 is signified as ****. **(d)** Depiction of scatter plots to compare the mean H-scores of β-catenin and FOXM1 in normal and colon carcinoma tissue samples, respectively. **(e)** Spearman’s rank correlation coefficient (r_s_) was depicted graphically by using the mean H-scores of β-catenin and FOXM1 from both normal and colon carcinoma tissues. **(f)** Multiple colon cancer cell lines (HCT 116 and HT29) were grown in 60 mm cell culture plates and treated with 20mM LiCl for 6 hours. Immunoblotting was done from whole cell lysates and probed against proteins of interest (FOXM1, β-catenin, Cyclin D1 as positive control, and Actin as loading control). **(g)** Whole-cell lysates obtained from HCT 116 and HT 29 cells after treatment with 20uM IWP2 inhibitor for 24 hrs, were used for immunoblotting against FOXM1 and β-catenin antibodies. **(h)** HCT 116 cell line was treated with purified Wnt (150ng/mL) for 24hrs. Immunoblotting was done against the respective antibodies in the figure. **(i)** HCT 116 and HT29 cell lines were transfected with 4 μg of pGZ-β-catenin or the empty PGZ vector and whole cell lysates were obtained after harvesting the cells post 24hrs from the time of transfection. Immunoblot depicts the expression of β-catenin, FoxM1, Cyclin D1, and Actin proteins. Densitometric analyses relative to Actin (loading control) are mentioned below the blots. **(j)** Using a converse approach, HCT 116 and SW480 cells, seeded in 60mm dishes, were transfected with 4 μg of either scramble shRNA or β-catenin shRNA plasmids and harvested post 48hrs from the experiment. Whole-cell lysates were obtained and immunoblotting was performed. Densitometry values are mentioned below the blots after comparison with a loading control. **(k)** HCT 116 cells were transfected with increasing concentration of PGZ-β-catenin plasmids (2 μg and 4 μg of DNA) and harvested for immunoblotting against selected antibodies as depicted in the figure. **(l)** cDNA prepared from PGZ-β-catenin and PGZ-transfected multiple colon cancer cell lines (HCT 116 and HT 29) were utilized in qRT-PCR analysis to compare and contrast the mRNA levels of genes of interest; FOXM1, β-catenin, and Cyclin D1. **(m)** In a converse approach, mRNA levels of β-catenin, FOXM1, and Cyclin D1 genes were observed through qRT-PCR analysis after preparing cDNA from 35mm plates of multiple colon cancer cell lines transfected with 2 μg of either scramble shRNA or β-catenin shRNA plasmids and harvested after 48hrs from time of transfection. Cyclin D1 (CCND1) served as positive control whereas 18S rRNA was used for normalization. The mean (+/-) s.d is represented by error bars; the p-values were calculated using independent, two-tailed Student’s t-test and p < 0.0001 is signified as ****.

We employed different measures to investigate the involvement of canonical Wnt signaling. Firstly, activation of the canonical Wnt signaling pathway can be attained through lithium chloride (LiCl), an inhibitor of glycogen synthase kinase-3β (GSK-3β). GSK-3β phosphorylates β-catenin in the cytoplasm and upon inhibition makes β-catenin free to translocate into the nucleus for signaling^30,31^. Colon cancer cell lines (HCT 116 and HT-29) were treated with 20mM LiCl for 6 hours. Immunoblots reveal an augmented increase in the levels of β-catenin, FoxM1, and Cyclin D1 proteins (Fig. 3f) suggesting the suspected involvement of Wnt signaling in regulating FoxM1. Secondly, IWP2, a small molecule inhibitor of Porcupine production, can inhibit the canonical and non-canonical Wnt signaling pathways^32^. So, HCT 116 and HT-29 cells were treated with 20uM IWP2 inhibitor and a significant decrease in the expression profiles of FoxM1 and Cyclin D1 was obtained (Fig. 3g). Thirdly, Wnt3a is a conserved, secreted glycoprotein that influences stabilization and nuclear translocation of β-catenin. Thus, purified Wnt 3a (150ng/mL) treatment was done in HCT 116 cell line for 24hours^33,34^. It led to substantial upregulation of FoxM1 protein levels with Cyclin D1 being the positive control^35^ (Figure 3h). These results suggest a strong correlation between Wnt signaling and FOXM1 regulation.

At the protein levels, β-catenin was overexpressed in multiple colon cancer cell lines (HCT 116, SW-480, and HT-29) and found to enhance FoxM1 protein expression. Cyclin D1, the positive control for β-catenin overexpression, displayed similar trend (Figure 3i). Conversely, the endogenous levels of β-catenin were depleted using short hairpin RNA (shRNA) and caused a stark reduction of FoxM1 expression (Figure 3j). Upon decreasing β-catenin in a dose-dependent manner in the HCT 116 cell line, a dose-dependent reduction in the expression of FoxM1 and Cyclin D1 were observed (Figure 3k).

To establish the influence of β-catenin at the transcript level, in multiple colon cancer cell lines (HCT 116, SW-480, and HT-29), β-catenin mRNA levels were overexpressed, that led to increased mRNA expression levels of the FOXM1 gene (Figure 3l and Supplementary Fig. S2). Conversely, β-catenin knockdown significantly reduced FoxM1 mRNA expression (Fig. 3m). Cyclin D1 served as the positive control. Altogether, these results establish the role of Wnt/β-catenin in mediating the regulation of the FOXM1 gene.

### p68 and β-catenin alliance controls expression of FOXM1 and its downstream targets in CRC

We sought to investigate the possible involvement of p68 and β-catenin towards the key downstream targets of the oncogenic transcription factor FOXM1^32^. IHC analysis of Survivin, downstream effector of FOXM1, was performed and observed increased expression of Survivin protein in colon carcinoma tissue samples compared to normal colon tissue (Fig. 4a). Quantification by H-scoring and the difference in staining intensities, represented through Notched Box Plot, between normal and carcinoma samples, were statistically significant for Survivin (Fig. 4b). Comparison of combined average H-scores of FOXM1 and Survivin, between normal and colon carcinoma samples showed increased levels in colon carcinoma compared to normal (Fig. 4c). The mean H-scores of Survivin was found to corroborate with the trends followed by p68, β-catenin, and FOXM1 in both normal and carcinoma samples (Fig. 4d). Upon using Spearman’s rank correlation test, H-scores of FOXM1 were found to bear strong positive correlation with H-scores of Survivin (r_s_ = 1.043). Similarly, there was a strong positive correlation between p68 and Survivin (r_s_ = 0.9886) (Fig. 4e). We also showed the comparative H-score profile of β-catenin with Survivin in both normal and colon carcinoma and found similar trends (Supplementary Fig. 3a). We have also illustrated a strong positive correlation between β-catenin and Survivin (r_s_ = 0.9321) based upon their H-scores and Spearman’s rank correlation test. Combined average H-scores of p68, β-catenin, and Survivin in normal and colon carcinoma were plotted and showed similar trends (Supplementary Fig. 3b). Consequently, the results reveal a highly plausible interconnection between p68 and β-catenin with Survivin.

**Fig 4.**
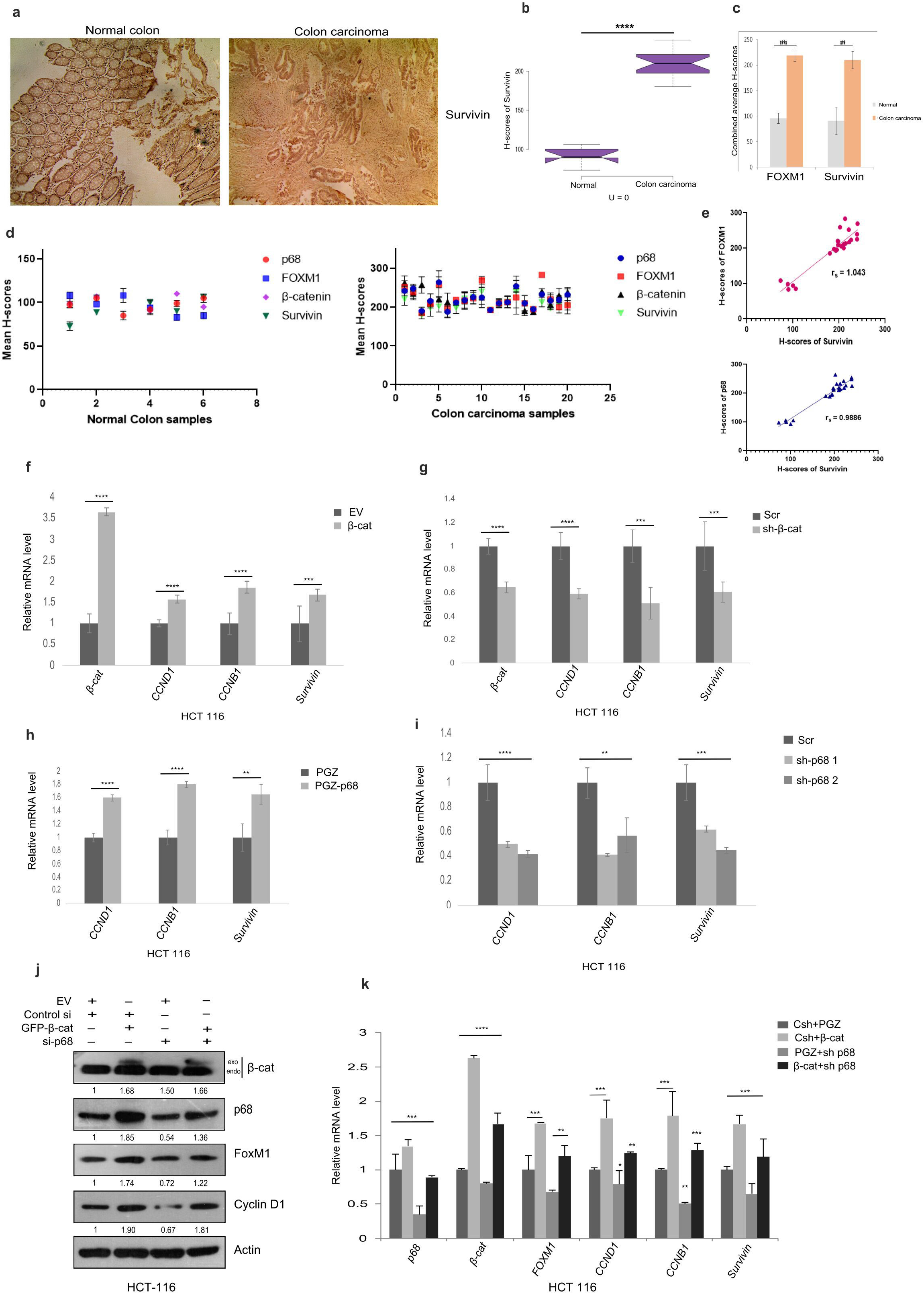
p68/ β-catenin synergism and their effects on downstream target genes of FOXM1. **(a)** Representative images obtained from Immunohistochemical (IHC) staining Survivin in human normal colon and colon carcinoma tissue samples. The images were captured at 200X magnification. **(b)** Representative Notched-box plots were obtained from the H-scores of Survivin in normal (n = 6) and colon carcinoma (n = 20) samples; H-scores were used for the calculation of Mann–Whitney U-values. **(c)** Comparison between the combined average H-scores of Survivin and FOXM1 in normal vs colon carcinoma samples. The mean (+/-) s.d is represented by error bars; the p-values were calculated using an independent, two-tailed Student’s t-test, and p < 0.0001 is signified as ****. **(d)** Depiction of scatter plots to compare the mean H-scores of p68, β-catenin, FOXM1, and Survivin in normal and colon carcinoma tissue samples, respectively. **(e)** Spearman’s rank correlation coefficient (r_s_) was depicted for the correlation between the mean H-scores of Survivin and FOXM1 along with the correlation between Survivin and p68 from both normal and colon carcinoma tissues. **(f)** to **(i)** HCT 116 cell line, plated in 35mm plates, was transfected with 2 μg each of PGZ-β-catenin, shRNA-β-catenin, PGZ-p68, and shRNA-p68 respectively along with the corresponding empty vector plasmids. They were harvested after 48 hrs and used for cDNA synthesis that was subsequently used for qRT-PCR analysis against the genes of interest (β-catenin, p68, FOXM1, Survivin, Cyclin B1, and Cyclin D1) **(j)** Rescue experiments were performed in HCT 116 cells. They were seeded in 60 mm plates and were transfected with 2 μg of scr or p68 siRNA, in combination with 2 μg of pGZ-β-catenin or EV. Whole-cell lysates were prepared after 48 hrs of experiment and probed for indicated proteins. Densitometry values below the blots are relative to the loading control (Actin). **(k)** Rescue experiments were performed in HCT 116 cells and probed for changes at the mRNA levels. They were seeded in 35 mm plates and were transfected with 1 μg of scr or p68 shRNA, in combination with 1 μg of pGZ-β-catenin or EV. The cells were harvested after 48hrs of experiment and cDNA was prepared that was subsequently used for qRT-PCR analysis against the indicated genes of interest. The mean (+/-) s.d is represented by error bars; the p-values were calculated using an independent, two-tailed Student’s t-test, and p < 0.0001 is signified as ****.

To further exemplify our findings at the transcript level, we checked the expression of target genes of FOXM1 *viz*., Survivin (responsible for enhanced survival) and Cyclin B1 (cell cycle regulator) through qRT-PCR. In HCT 116 cells, it was observed that the increase/decrease in cellular β-catenin caused similar trend (augmented/decreased levels) of Cyclin B1 and Survivin (Fig. 4f and 4g). Similarly, overexpression and downregulation of p68 led to elevated and decreased levels of Survivin and Cyclin B1 at the transcript levels respectively (Fig. 4h and 4i). Cyclin D1 served as the positive control. These findings exemplify the influence of p68 and β-catenin on FOXM1 target gene expression and establishes a strong foothold of these molecular players on the regulation of FOXM1 gene expression.

Through a combinatorial approach, in HCT 116 cells, it was proved through Immunoblot and qRT-PCR experiments that β-catenin-mediated increase in the protein and mRNA levels of FOXM1 and its signature target genes (Survivin and Cyclin B1) got abrogated under p68-depleted conditions (Fig. 4j and 4k). This highlights that p68 and β-catenin collaboration is required for FOXM1 expression. Cyclin D1 served as a positive control for both p68 and β-catenin signaling. Collectively, the above results highlight a strong impact of p68/β-catenin interplay and corroborate that p68 along with β-catenin exerts a strong influence on expression of FOXM1 and its downstream effector target genes.

### p68 works in alliance with β-catenin to regulate the FOXM1 gene expression

To gain deeper insights, we speculated that p68 might be regulating the promoter activity of FOXM1 by transcriptionally coactivating it by allying with β-catenin^36,37^. We considered the FOXM1 promoter region of ∼1.3 kb upstream to the translation start site (TSS) and found that it contains six putative β-catenin/TCF-LEF-binding elements (TBEs).

Chromatin Immunoprecipitation (ChIP) assay was conducted to decipher the cooperative synergism of p68 and β-catenin in regulating FOXM1. The schematic representation of the FOXM1 promoter is given for reference (Fig. 5a). ChIPs were carried out with sonicated crosslinked chromatin isolated from treated HCT 116 cells using anti-p68, anti-β-catenin, and anti-TCF4 antibodies. It was followed by amplification of different TBE (TBE 1-2 and TBE 3-6) sites on the FOXM1 promoter. ChIP data (Fig. 5b) confirmed the *in vivo* occupancy of the TCF/LEF sites on FOXM1 promoter by p68, β-catenin, and TCF4. Amplification of Cyclin D1 promoter region containing TCF sites served as the positive control. GAPDH aided as the negative control. Consequently, we concluded that p68, in alliance with β-catenin regulates FOXM1 by physically occupying its TCF4 sites.

**Fig 5.**
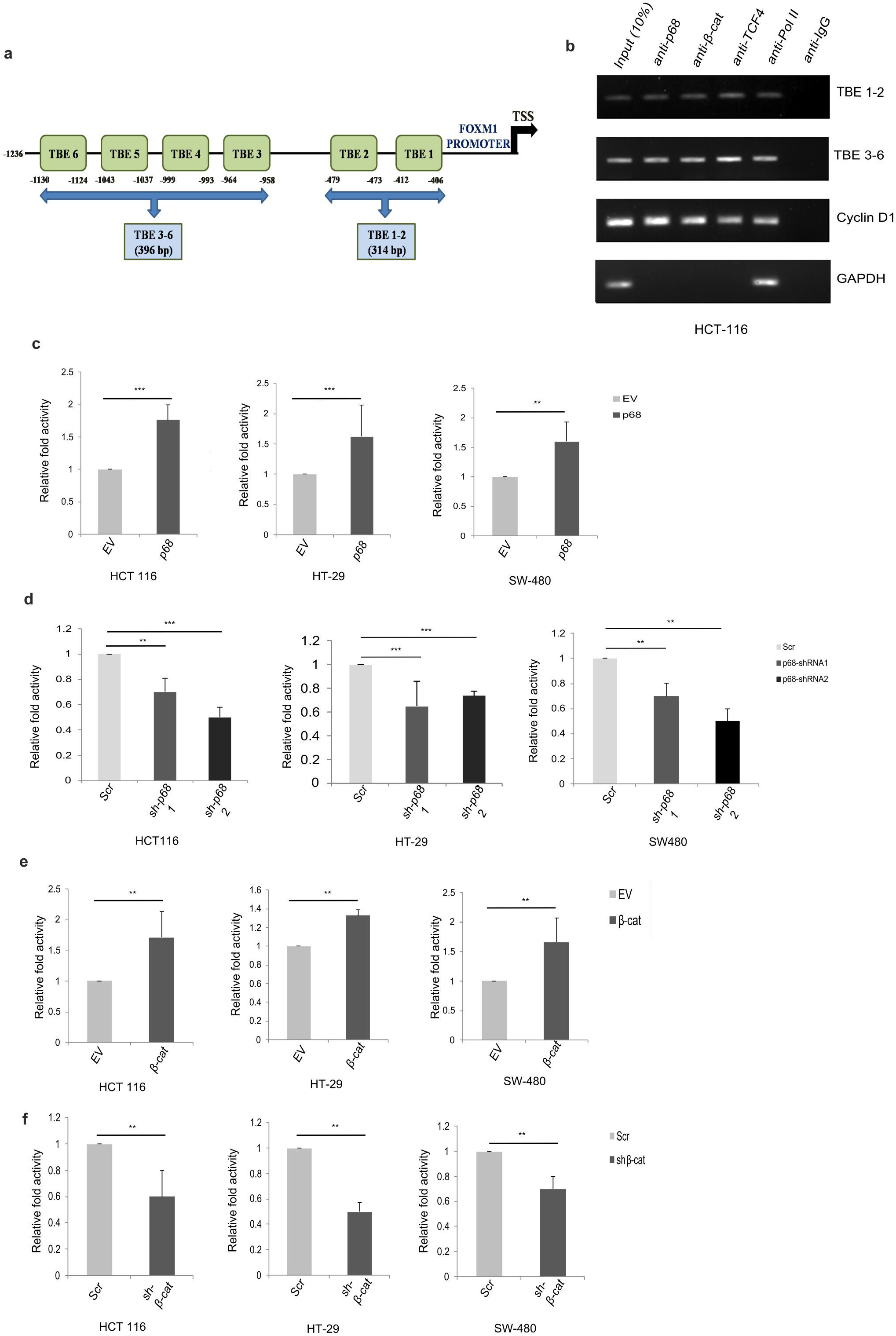
p68 works in alliance with β-catenin to transcriptionally regulate the FOXM1 gene through influencing FOXM1 promoter activity. **(a)** Schematic representation of the 1.3kb cloned human FOXM1 promoter displaying multiple TCF/LEF binding elements (TBE) sites. **(b)** HCT 116 cells were grown in 100 mm culture dishes. They were harvested for endogenous Chromatin immunoprecipitation (ChIP) assay using the indicated antibodies. DNA extract (10% without ChIP) was used as input. The Cyclin D1 promoter region was chosen as the positive control for both p68 and β-catenin. Actin promoter was the positive control for Pol II. IgG was the negative control of the experiment. The immunoprecipitated DNAs were PCR amplified using primers designed from the FOXM1 promoter region named TBE 1–2 (314 bp), flanking sites TBE1 and 2 along with TBE 3-6 (396 bp), flanking TBE 3, 4, 5, 6. **(c)** All the three colon cancer cell lines (HCT 116, SW 480, HT29) were seeded in 35 mm culture dishes. They were transfected with either 1 μg pIRES-p68 or pIRES-EV along with pGL3-WT-FOXM1-promoter construct (1 μg) and Renilla luciferase plasmid (50 ng). Cells co-transfected with EV (pIRES), pGL3-WT-FOXM1-prom and Renilla luciferase plasmid served as control. The luciferase activity was measured 48hrs post transfection. **(d)** All the three colon cancer cell lines (HCT 116, SW 480, HT29) were seeded in 35 mm culture dishes. They were transfected with either 1 μg shRNA-p681 and 2 or scrambled shRNA (EV) together with pGL3-WT-FOXM1-promoter construct (1 μg) and Renilla luciferase plasmid (50 ng). The luciferase activity was measured 48hrs post-transfection. **(e), (f)** Luciferase assay was conducted in all the three colon cancer cell lines (HCT 116, SW 480, HT29) after transfecting with pGZ-β-catenin or EV; or shRNA-β-catenin or Scrambled along with pGL3-WT-FOXM1-promoter construct (1 μg) and Renilla luciferase plasmid (50 ng). Renilla luciferase was used for normalization and data is characterized in terms of fold change activity concerning control. Error bars in all the indicated sub-figures represent mean (+/-) s.d from three independent biological repeats and the p-values were calculated using an independent, two-tailed Student’s t-test where p < 0.0001 is signified as ****.

A luciferase assay is a very sensitive and convenient way to examine the transcriptional activity of a gene^38^. To explore the mechanistic perceptions of novel regulation of the FOXM1 gene, 1236 bp fragment of the human FOXM1 promoter was successfully cloned in PGL3 basic vector through conventional cloning procedure used previously. Next, the luciferase assays were conducted in multiple colon cancer cell lines (HCT 116, SW-480, HT-29). Results demonstrated that p68 overexpression, significantly upregulated FOXM1 promoter activity (Fig. 5c). Conversely, the knockdown of p68 through multiple shRNAs in all three colon cancer cell lines (Fig. 5d) and siRNA in HCT 116 cell line (Supplementary Fig S4) showed a prominent decrease in the FOXM1 promoter activity. Similar results were obtained upon overexpressing β-catenin and concomitant downregulating it by using specific shRNA (Fig. 5e and 5f). Therefore, both p68 and β-catenin are involved in the transcriptional regulation of the FOXM1 promoter and regulates the elevation and diminution of its promoter activity under overexpressed and knocked-down scenario respectively.

### Detailed mechanistic characterization of p68/β-catenin synergism on FOXM1 promoter

To mechanistically explicate the regulatory axis, we embraced a combinatorial approach to establish the synergistic effect of p68 and β-catenin on the FOXM1 promoter activity. p68 and β-catenin were overexpressed either individually or in a combinatorial manner to observe their synergistic effect. The combinatorial luciferase experiments were carried out in HCT 116, SW-480, and HT-29 cell lines. It yielded positive results with an enhanced cooperative synergistic effect of both p68 and β-catenin leading to an intensification of FOXM1 promoter activity (Fig. 6a). Additionally, by rescue experiments, the role of p68 was profoundly observed, whereby the increased promoter activity under β-catenin overexpressed condition was diminished upon reducing p68 level through shRNA. Furthermore, under the combined scenario of β-catenin overexpression and p68 knockdown state, the promoter activity of FOXM1 was rescued. Overall, these results strengthen our hypothesis of p68 coactivating β-catenin in mediating the transcriptional activation of the FoxM1 promoter (Fig. 6b).

**Fig 6.**
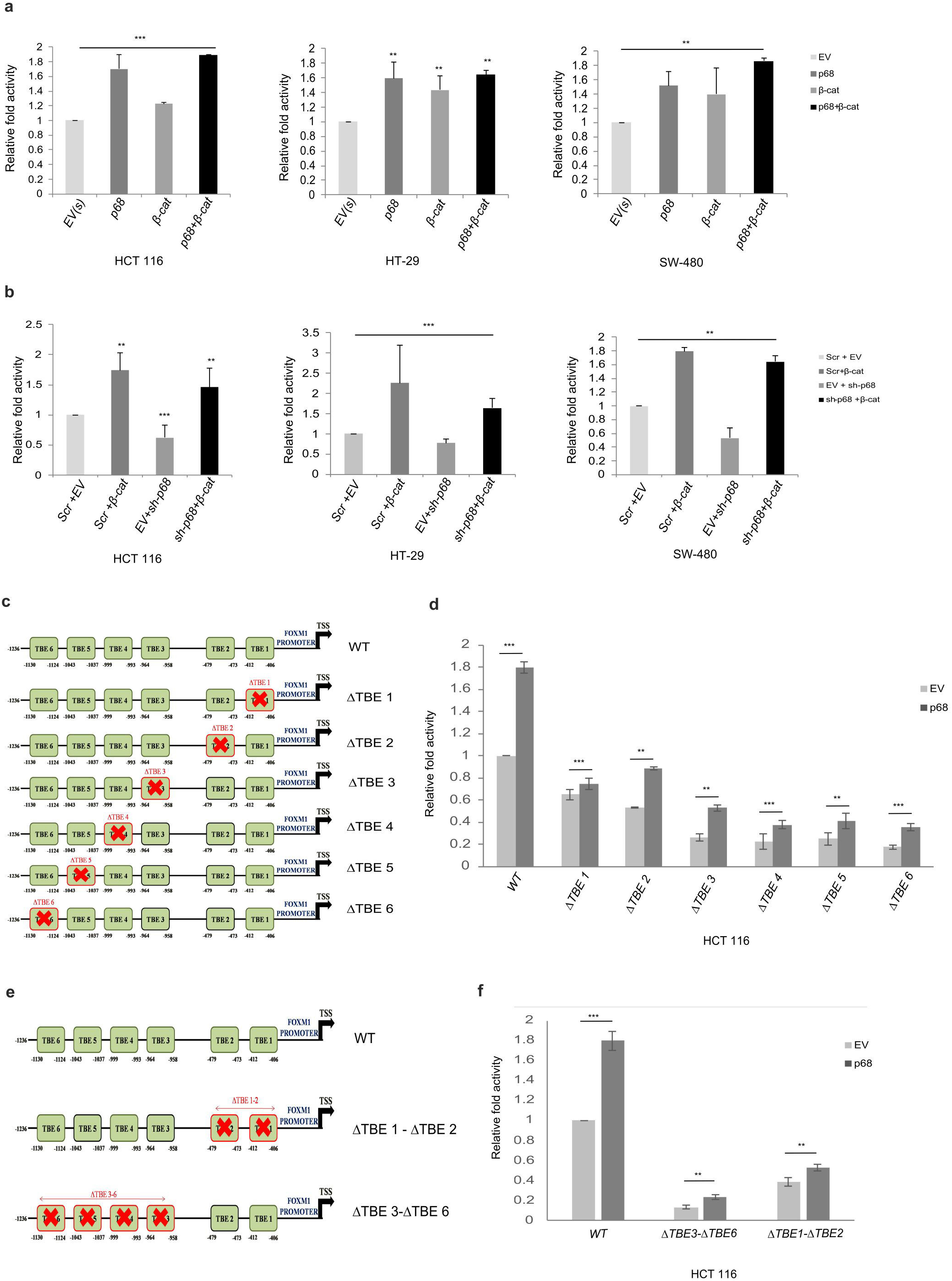
Mechanistic elucidation of p68/β-catenin synergism on FOXM1 promoter activity effect on regulation through mutation of FOXM1 promoter. **(a)** Combinatorial dual luciferase assay was conducted on all three colon cancer cell lines (HCT 116, SW480, HT29). The cells were seeded in 35 mm culture dishes and transfected with either EV, PGZ-p68, PGZ-β-catenin, and a combination of both PGZ-p68 and PGZ-β-catenin along with an equivalent concentration of pGL3-WT-FOXM1-promoter construct and Renilla luciferase plasmid (50 ng for normalization). The luciferase activity was measured 48hrs post-transfection. **(b)** Rescue experiments for defining the supremacy of p68 were used for luciferase measurements. In all the three colon cancer cell lines (HCT 116, SW480, HT29), transfection was performed with either EV or PGZ-β-catenin or shRNA-p68 or a combination of both shRNA-p68 and PGZ-β-catenin along with an equivalent concentration of pGL3-WT-FOXM1-promoter construct and Renilla luciferase plasmid (50 ng for normalization). The luciferase activity was measured 48hrs post-transfection **(c)** and **(e)** Schematic representation of Site-directed mutagenesis conducted upon the 6 TBE sites on the 1.3kb cloned human FOXM1 promoter. Deletion mutants were constructed using Site-directed mutagenesis (SDM) where each of the six TCF4 sites was deleted individually and designated as ΔTBE1 to 6 respectively. Also combinatorial multiple site deletion approaches were employed by deleting both TBE1 and TBE2 in one construct (ΔTBE1-2) and the entire flank of TBE3 to TBE6 in another construct (ΔTBE3-6). **(d)** and **(f)** Promoter activity measurement was measured using luciferase assay which was conducted to compare and contrast using pGL3-WT-FOXM1-promoter or pGL3-mutant-FOXM1-promoters (ΔTBE1 to 6) along with transfecting with an equivalent concentration of pGL3-WT-FOXM1-promoter construct and Renilla luciferase plasmid (50 ng for normalization). The luciferase activity was measured 48hrs post-transfection. Promoter activity was also measured by transfecting the plates with the mutant constructs in the form of ΔTBE1 to ΔTBE6 mutant FOXM1 promoter groups. The luciferase activity was measured 48hrs post-transfection. Error bars denote mean (+/-) s.d from three independent biological repeats. Indicated p-values were calculated using Student’s t-test and p < 0.0001 is represented as **** (highly significant).

Next, to exclusively delineate the role of p68 in regulating FOXM1, we sought to explore the promoter activity of FOXM1 by mutating the putative TBE sites. The wild-type promoter of FOXM1 contained 6 TCF4 binding sites referred to as TBE and were designated as site TBE1 – CTTTGTA (-406 to -412), TBE2 GCCAAAG (-473 to -479), TBE3 – CTTTGTA (-958 to -964), TBE4 – CTTTGCA (-993 to -999), TBE5 – CTTTGTA (-1037 to -1043), and TBE6 – CTCAAAG (-1124 to -1130). The positions were in alignment and relative to the transcription start site. Deletion mutants were constructed using Site-directed mutagenesis (SDM) whereby each of the six TCF4 sites was deleted individually and designated as ΔTBE1, ΔTBE2, ΔTBE3, ΔTBE4, ΔTBE5, and ΔTBE6 respectively (Fig. 6c). Moreover, deletion of multiple sites in combination was done wherein both TBE1 and TBE2 were deleted (ΔTBE1-2) and the entire flank of TBE3 to TBE6 was deleted (ΔTBE3-6) (Fig. 6e). Upon p68 overexpression in HCT 116 cells, the promoter activity of FOXM1 revealed a substantial reduction in case of deletion mutants compared to the wild type (Fig. 6d). Besides, luciferase assays in the multiple deletion constructs (TBE1-2 and TBE3-6) revealed highly diminished promoter activity profiles of mutant FOXM1 promoter compared to wild type promoter under p68 overexpressed conditions. Interestingly, there was a huge abrogation of the promoter activity of FOXM1, when deletion mutant ΔTBE3-6 was used (Fig. 6f). Thus, we can mechanistically and coherently conclude that p68 exerts its function through β-catenin/TCF4 complex in the transcriptional activation of the FOXM1 promoter at the TBE sites. Upon mutating these sites, an abrogation of transcriptional activity of the FOXM1 promoter was observed.

### p68-dependent enhanced proliferation and migration of CRC cells get inhibited upon FOXM1 inhibition -indicating involvement of “p68-FOXM1” axis

Wound healing assay was performed after overexpression of p68 and β-catenin separately in HCT 116 and observing its influence on the wound created at time intervals of 0, 24 and 48 hrs. It was observed that an increased percentage of wound healing (decreased wound area), through accelerated gap filling, occurred due to overexpression of either p68 or β-catenin (Fig. 7a). This implicates their involvement in cellular proliferation and migration. Using a converse approach, both p68 and β-catenin were downregulated using respective shRNAs, and it was observed that there wasn’t any significant change in the percentage of wound healing implying diminishing proliferative rates of cells (Supplementary Fig S5). Also, LiCl and Wnt 3a treatment were administered and the wound was observed at the indicated time points. Consequently, an increased percentage of healing of the wound and decreased area after 24 hours of the treatment was observed, thereby indicating that Wnt3a-mediated activation of canonical Wnt/β-catenin signaling aids in cellular proliferation (Fig. 7b and c).

**Fig 7.**
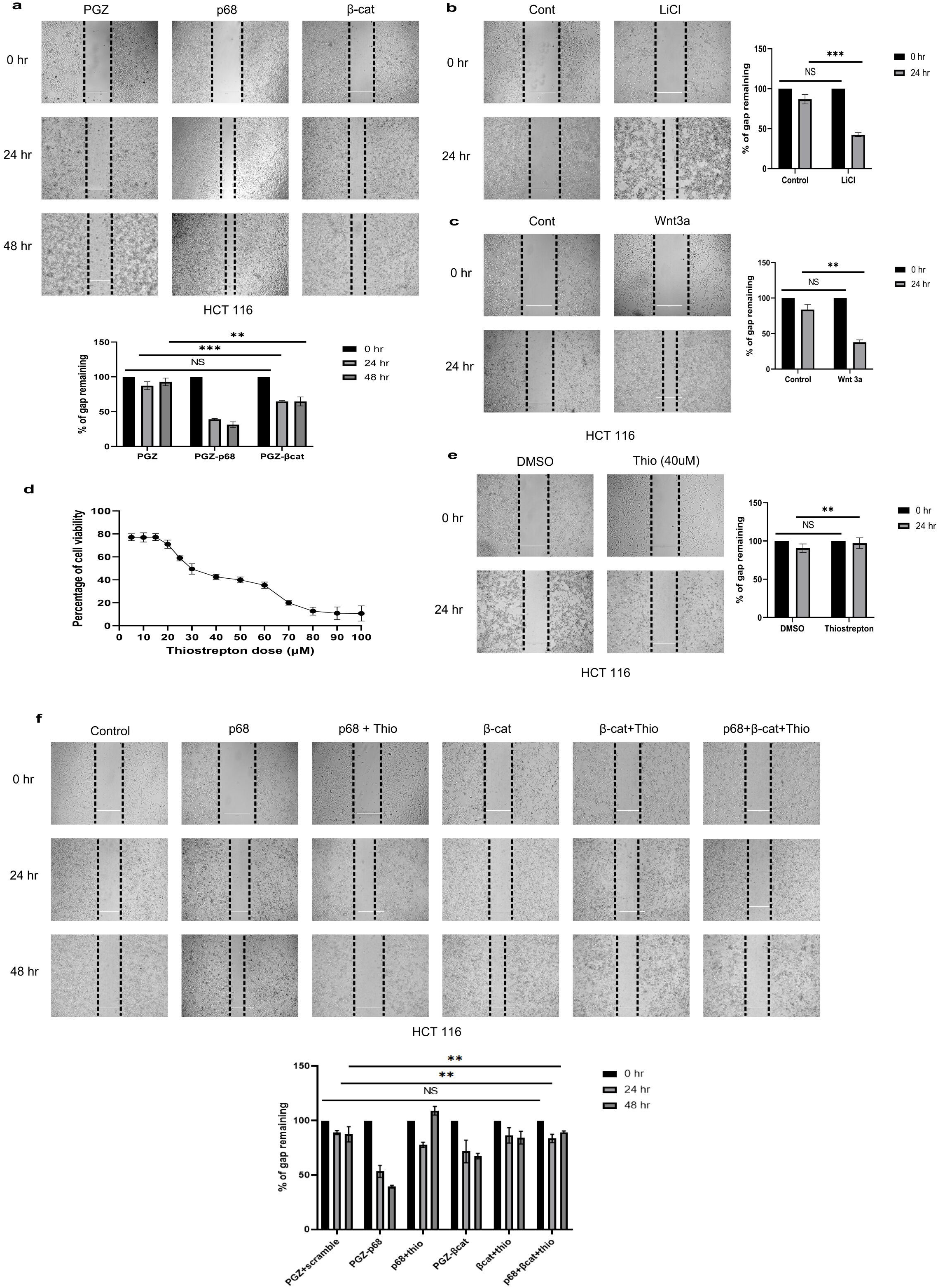
Heightened cellular proliferation and migration due to p68-mediated regulation of FOXM1 and treatment with FOXM1 inhibitor Thiostrepton. **(a)** Wound healing assay was conducted in HCT 116 cells along with experimental transfections. The cells were seeded in 35 mm cell culture dishes and were treated with EV or PGZ-β-catenin or PGZ-p68. 10ul sterile pipette tips were used to make transverse scratches on the plates and the images were captured at the specified time points. Scale bar-400um. Bar graph signifies the percentage of gap filling. E, HCT 116 cells seeded in 35 mm cell culture dishes were transfected with 2ug of EV or CHIP. After making transverse scratches on the plates with 10ul sterile pipette tips, the images were captured at the specified time points. Scale bar-400um. **(b), (c), (e)** The cells were treated with LiCl (6hrs) or Wnt3a (24hrs) or Thiostrepton (24hrs) along with DMSO or water as empty control, and the induced scratch was observed at intervals of 0 hr, 24 hr, 48 hrs. **(d)** HCT 116 cells treated with DMSO or Thiostrepton (24 hours) were checked for their cell viability properties through MTT assay. The average absorbance values of three biological repeats were plotted. **(f)** A multi-sectoral combinatorial approach was employed. HCT 116 cells were seeded in 35 mm culture dishes and thereafter transfected with EV or PGZ-p68 or PGZ-p68 with the treatment of thiostrepton or PGZ-β-catenin or β-catenin with thiostrepton treatment, or combination of p68, β-catenin, and treatment with thiostrepton. Scratch assay was performed and graphical bar diagrams represent the percentage of gap filling due to reduction/increase of the area of the wound. Error bars in all the indicated subfigures represent mean (+/-) s.d. from three independent biological repeats. Indicated p values were calculated using Student’s t-test and p≤0.0000 is represented as ****.

Furthermore, we thought of using an established FOXM1 inhibitor, Thiostrepton to comprehensively dissect the oncogenic potential of FOXM1^39^. The upregulation of FOXM1 mediated by p68 subsequently predisposes cells to tumorigenesis and leads to enhanced cellular proliferation and migration. To combat such unregulated cell growth, the influence of FOXM1 inhibitor needed to be deciphered. Firstly, an MTT assay was conducted in HCT 116 cells by treating with sequential doses of thiostrepton (5, 10, 15, 20, 25, 30, 40, 50, 60, 70, 80, 90, 100μM) for 24hrs. It was revealed that with increasing doses, the viability of HCT 116 cells kept on decreasing (Fig. 7d). Next, a wound healing assay was performed in HCT 116 cell line after treatment with 40μM thiostrepton. It was observed that the proliferative influence of FOXM1 was significantly diminished through this inhibitor (Fig. 7e). Next, we employed a combinatorial approach for dissecting the ‘p68-β-catenin-FOXM1’ axis by using thiostrepton to decelerate the oncogenic pace. Wound healing assay was conducted at 0, 24, and 48 hrs intervals by utilizing the following combinatorial approaches: (a) overexpressing p68with and without thiostrepton, (b) overexpressing β-catenin with and without thiostrepton, and (c) overexpressing both p68 and β-catenin, along with the thiostrepton treatment in HCT 116 cells. Interestingly, the p68-mediated cell proliferation and migration attributes get abrogated upon FOXM1 inhibition. Inhibition of FOXM1 by thiostrepton significantly decreased cell proliferation and wound healing that was otherwise pronounced under p68 and β-catenin overexpression conditions (Fig. 7f). Therefore, we successfully established the ‘p68-β-catenin-FOXM1’ axis in oncogenesis and the importance of FOXM1 inhibition as future avenues for CRC treatment.

### Effect of the ‘p68/β-catenin/FOXM1’ axis in colon carcinogenesis

FOXM1 is a cell cycle regulator and increases the S-phase population^40^. Cell cycle analysis after treating the cells with thiostrepton for 24hrs revealed an inhibitory action (Supplementary Figure S6a). A significant rise in the percentage of S-phase cells was observed upon overexpression of both p68 and β-catenin together. However, the percentage of S-phase cells declined upon treatment with thiostrepton under a similar condition. Decrease in the percentage of S-phase cells was observed in p68-β-catenin-thiostrepton combination treatment (Fig. 8a). Furthermore, colony formation assay disclosed the vigorous colony forming ability of HCT 116 cells under overexpressed p68 and β-catenin conditions (Fig. 8b). Next, we treated these cells with LiCl and observed an enhanced colony formation (Supplementary figure S6b). Utilizing a converse approach, upon inhibition of p68 expression using concomitant siRNA treatment against it and downregulation of β-catenin through shRNA-mediated knockdown, a drastic reduction in the number of colonies was observed (Fig. 8c and Supplementary figure S6c). These results clearly indicate the importance of p68-mediated regulation *via* canonical Wnt/β-catenin signaling in enhancing cellular proliferation, migration, cell cycle profiles, and colony formation abilities. To highlight the importance of FOXM1, we inhibited it in a dose-dependent manner by using 20, 40, and 60 μM of thiostrepton and investigating its effect on clonogenic capabilities in HCT 116 cells. It was found that colony formation was significantly inhibited by thiostrepton and found to be extremely diminished in 60 μM (Fig. 8d). Thereafter, we employed the combinatorial approach as depicted in the figure and found diminishing profiles of colony formation under over-expressed p68 and β-catenin conditions when treated with thiostrepton in comparison with the untreated controls (Fig. 8e).

**Fig 8.**
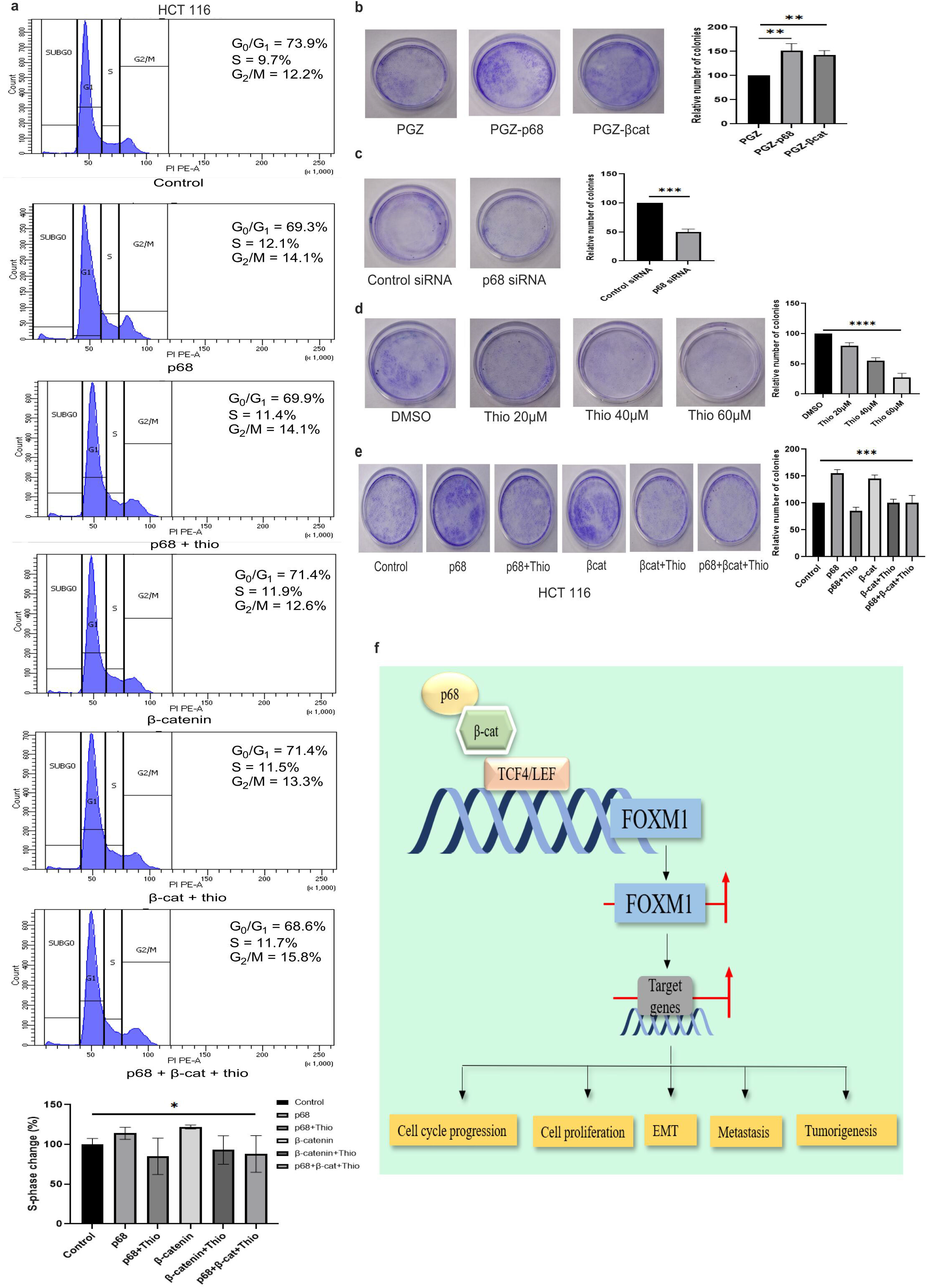
Increased hallmarks of cell cycle and colony forming features due to p68-mediated regulation of FOXM1 and support to oncogenesis. **(a)** Cell cycle distribution profiles of HCT 116 cells transfected with EV or PGZ-p68 or PGZ-p68 with the treatment of thiostrepton or PGZ-β-catenin or β-catenin with thiostrepton treatment, or a combination of p68, β-catenin, and treatment with thiostrepton was done carefully for subsequent flow cytometry. The graph signifies the change in the percentage of S phase cells in the treated compared to the control. **(b)** HCT 116 (5000 in number) cells were seeded in 35mm culture dishes and made stabilized before transfection (EV. PGZ-p68 and PGZ-β-catenin) and monitored regularly for their colony formation features. After 2 weeks, the plates were stained with crystal violet, washed, and observed. **(c)** HCT 116 cells were transfected with control siRNA and p68 siRNA and harvested according to the colony formation protocol. The graphs represent the percentage of colony formation **(d)** Colony formation assay was done in HCT 116 cells after treatment with increasing doses (20, 40, and 60 μM) of thiostrepton. **(e)** HCT 116 cells were transfected with the combination of EV or PGZ-p68 or PGZ-p68 with the treatment of thiostrepton or PGZ-β-catenin or β-catenin with thiostrepton treatment, or a combination of p68, β-catenin, and treatment with thiostrepton for subsequent flow. The relative number of colonies was represented in graphs and averaged from three biological repeats. **(f)** Schematic representation of the novel mode of regulatory axis involving gene regulation of FOXM1 via p68 coactivation mediated by Wnt/β-catenin signaling in potentiating colon carcinogenesis. The mean (+/-) s.d is represented by error bars; the p-values were calculated using an independent, two-tailed Student’s t-test, and p < 0.0001 is signified as ****.

In summary, our work presents compelling pieces of evidence for potentiation of colon carcinogenesis through FOXM1 and mechanistically characterizing its regulation by the master regulator p68 through the coactivation of the canonical Wnt/β-catenin signaling pathway. The schematic diagram summarizes the regulation of FOXM1 by p68 through coactivation of β-catenin and their occupancy on the TCF/LEF sites of the FOXM1 promoter to elicit enhanced transcription. Increased FOXM1 expression subsequently mediates its downstream impact on the colon carcinogenesis hallmarks (Fig. 8e).

## Discussion

The hallmarks of cancer in terms of cellular proliferation, migration, clonogenic potential, evasion of apoptosis, and metastasis threaten the normal cellular homeostasis^41^. Despite improvements in detection, diagnostic and therapeutic avenues, the prognosis of CRC still shows dismal rates^9^. Thus, a comprehensive analysis of signaling cross-talks and molecular players that trigger its initiation and progression is required. FOXM1 is found to be aberrantly overexpressed in multiple cancers through higher rates of proliferation, metastasis, influences tumor initiation and progression^21^. Furthermore, FOXM1 has been hailed as a potential molecular marker in CRC^15,19^. However, reports on its gene regulation aspect are limited. Therefore, articulate understanding of mechanistic complexities behind the regulation of FOXM1 may help in the effective clinical management of CRC. p68 is highly implicated in most cancers and has been established as a master regulator because of its multifaceted functionalities ranging from RNA processing and modification to transcriptional coactivation potential. The previous studies reported from our laboratory have strengthened its role as a coactivator in the oncogenic scenario ^36,42,43,44^.

In this study, we sought to unravel a novel mode of regulation of FOXM1 by the coactivation potential of p68. We used bioinformatic analyses (UALCAN and TCGA) to explore the heightened expression profiles of p68 and FOXM1 in CRC. Through immunohistochemistry in normal and colon carcinoma patient tissue samples, it was revealed that there was a strong correlation between p68 and FOXM1. Subsequently, through immunoblotting and qRT-PCR analysis in multiple colon cancer cell lines, we found that enhancement of p68 expression led to elevated expression of FOXM1 and the converse stood true in case of downregulation. To examine the influence of p68 coactivation, we contemplated the importance of Wnt signaling due to several compelling evidences linking p68 with Wnt signaling and β-catenin^45,43^. Using IHC analyses and statistical data, we found a strong positive correlation between β-catenin and FOXM1 in colon cancer patient tissue samples compared to normal samples. To elaborate our studies, we utilized Wnt activators like LiCl and purified Wnt 3a ligand^30,31^. The expression of FoxM1 protein synced with these treatments and showed increased levels along with a concomitant increase of Cyclin D1, the positive control of Wnt signaling. Next, we overexpressed and knocked down the β-catenin level and found that FOXM1 showed a comparable increase and decrease at both protein and mRNA levels. To exclusively delineate the regulation of FOXM1, its downstream target genes needed to be considered. FOXM1 stimulates the expression of a variety of genes namely Aurora B, Cyclin B1, Plk1, CDC25B, CENP-A, and Survivin^46^. We conducted IHC studies on normal and colon carcinoma patient tissues to find the expression of the Survivin protein. It was found to correlate with the heightened expression profile of FOXM1 and showed a strong positive correlation with p68, β-catenin, and FOXM1. To exemplify our findings, mRNA profiles of FOXM1 downstream target genes, Survivin and Cyclin B1 were checked under conditions of overexpressed or knocked down p68 and β-catenin. Both the target genes increased or diminished according to the elevated or depleted levels of p68 and β-catenin. To highlight the overall importance of p68, rescue experiments were conducted, where an increase in the protein and mRNA levels of FOXM1 and its target genes (*viz*., Survivin and Cyclin B1) mediated by β-catenin was abrogated under p68-depleted conditions. Furthermore, we analyzed the promoter of FOXM1 gene using ALGGEN PROMO software and found the existence of crucial TCF/LEF binding sites on the FOXM1 promoter. After cloning 1.3kb region of the human FOXM1 promoter, we conducted a chromatin immunoprecipitation (CHIP) assay to examine the binding efficiency of p68 and β-catenin on the putative Wnt signaling activating TCF/LEF sites on the FOXM1 promoter. The results established the physical occupancy of p68 and β-catenin on the FOXM1 promoter. Thereafter, the strength of the FOXM1 promoter activity was measured using luciferase assay. Increased or decreased FOXM1 promoter activity was synchronized with the overexpression and knocked down of p68 and β-catenin. To strengthen our mechanistic studies and ascertain the influence of p68/β-catenin synergism, we overexpressed them singly and/or in combination and found heightened FOXM1 promoter activity in colon cancer cell lines. Next, we conducted rescue experiments to elicit the importance of p68 in the entire signaling scenario. Our novel findings were further corroborated with site-directed mutagenesis studies. Interestingly, under p68 overexpressed conditions, the FOXM1 mutated promoter showed a sharp decline in its activity compared to the wild-type promoter. Also, the promoter activity declined extremely upon expression of the mutant promoters having multiple TCF/LEF site mutations (ΔTBE1-2 and ΔTBE3-6) compared to wild-type promoter. To strengthen these studies, we further used functional assays to prove the oncogenic potential of the regulatory axis. Wound healing assays were conducted to establish cellular proliferation and migration. The percentage of wound healing increased upon activating the Wnt signaling and overexpressing p68. Thiostrepton, an inhibitor of FOXM1^39,47,48^, was used to fully characterize and dissect the functional assays and their importance. MTT assay showed a decreasing trend of cell viability with increasing doses of thiostrepton. We also designed a unique combinatorial experimental design where p68 and β-catenin were expressed either individually, conjointly, or in combination with thiostrepton drug to notice the effect of FOXM1 gene regulation. It was found that thiostrepton inhibited the increased manifestation of FOXM1 and abrogated the heightened levels of wound healing under the influence of p68. Finally, we conducted cell cycle analysis in a similar combinatorial approach which yielded positive results. Through functional assays, the colony-forming abilities of colon cancer cells under explicit stimulation by p68 and β-catenin were revealed. Through thiostrepton treatment, the clonogenic abilities diminished significantly. The depleting effect was strong even under p68 and β-catenin overexpressed conditions, either used alone or in combination. Thiostrepton, by inhibiting the elevated FOXM1 expression, in terms of regulating cellular proliferation, migration, cell cycle progression, and colony formation ability, unraveled the importance of FOXM1 in the oncogenic scenario.

Therefore, the current study brings forth a novel mode of regulation of FOXM1 by p68 acting through the coactivation of Wnt signaling and bringing forth complex crosstalk that cooperatively drives oncogenesis. Further exploration of the novel regulatory axis of ‘p68-β-catenin-FOXM1’ can steer the direction of basic and translational scientific research toward the successful management of colon carcinogenesis.

## Materials and methods

### Human colon tissue samples

Formalin-fixed paraffin-embedded (FFPE) post-surgical human normal colon (n = 6) and colon carcinoma (n = 20) tissue samples were collected following the clinical and ethical approval and regulations by the ethical committee of both CSIR-IICB and Park Clinic^49^.

### Cell culture, transfection, and chemicals

Human colorectal cancer (CRC) cell lines [HCT 116, SW-480, HT-29] and human embryonic kidney cell line [HEK-293] were procured from ATCC (Manassas, VA, USA) were cultured as described previously. Lipofectamine 2000 and 3000 (Invitrogen) were used for transfection experiments and Lipofectamine RNAi MAX (Invitrogen) for siRNA-based experiments. Thiostrepton (Sigma-Aldrich, Taufkirchen, Germany) was used in < 0.1% DMS.

### Expression plasmid constructs and cloning

Subcloning of p68 was done from the pSG5-Myc vector into pGZ21dx, pEGFP-c3, and pIRES-hrGFP-1a. Similarly, pBI-βcatenin was sub-cloned into pGZ21dx vector and the human Wnt3a gene was cloned in pcDNA3.1-myc-his, respectively in our laboratory. PCR amplification of the FOXM1 promoter region (1.3kb fragment) was done using genomic DNA from HEK-293 and cloned into pGL3 basic vector using standard procedures. ALGGEN-PROMO (http://alggen.lsi.upc.es/) was referred for analysis of putative binding sites. Restriction enzyme-based digestion verified all constructs. Final confirmation was provided by sequencing. All the primers were ordered from IDT (Integrated DNA Technologies, Coralville, IA, USA) and the sequences are provided in the Supplementary file 1.

### Site-directed mutagenesis

Deletion of TBE consensus sites in the FOXM1 promoter was accomplished using the QuickChange XL Site-Directed Mutagenesis^44^ kit (Agilent Technologies, Santa Clara, CA, USA) by following manufacturer’s instructions and was then cloned into the pGL3 basic vector. Primer sequences are elaborated in Supplementary file 1.

### RNA interference-mediated knockdown

Two different p68-shRNAs along with a control (scramble) shRNA were subcloned in pMKO.1-Puro plasmid. β-catenin shRNA was purchased from Addgene (pLKO.1 puro shRNA β-catenin #18803). Scramble siRNA and p68 siRNA were purchased from Santa Cruz Biotechnology (Santa Cruz, CA, USA) and used at a final concentration of 30 nM.

### Immunoblot analysis

The whole cell lysates (WCL) were prepared and immunoblot analyses were performed as described previously^43,50^. The primary antibodies that were used are p68 (Abcam); Cyclin D1, FOXM1, Survivin, TCF4, HRP-tagged anti-rabbit and anti-mouse secondary antibodies (Cell Signaling Technology); Actin, HRP-tagged anti-goat secondary antibody (Sigma-Aldrich), GFP, β-catenin, and GAPDH (Santacruz Biotechnology). Evaluation was done using Luminata Classico Western HRP Substrate (Millipore) following the manufacturer’s protocol. The computation of the densitometry values was done using GelQuant.Net software (http://biochemlabsolutions.com/GelQuantNET.html).

### Quantitative real-time PCR

Trizol reagent (Invitrogen, NY, USA) was used to extract total RNA and converted to cDNA using RevertAid H Minus First Strand cDNA Synthesis Kit (Thermo Scientific™). qRT-PCR analysis was done using FastStart Universal SYBR Green Master (Roche) on a ViiA 7 Real-Time PCR system (Applied Biosystems). 18S rRNA served as the internal control. The results and calculations were based upon three independent experiments in triplicate repeats. Primers are mentioned in Supplementary file 1.

### Luciferase-based promoter activity assay

pGL3-FOXM1 promoter luciferase reporter constructs and Renilla luciferase plasmid (pRL-TK) were used to perform Dual Luciferase Assay 48 hrs post-transfection. A minimum of three biological repeats and three technical repeats were used for proper quantification. Luciferase activity was determined luminometrically in the Varioskan Flash Multimode Reader (Thermo Fisher Scientific, Waltham, MA, USA). Dual luciferase assay system kit (Promega, Madison, WI, USA) was used following manufacturer’s protocol as described previously^51^.

### Chromatin Immunoprecipitation (ChIP) assay

ChIP was performed using indicated antibodies. The antibody against RNA Polymerase II (Pol II) and IgG served as a positive and negative control, respectively. The immunoprecipitated DNAs were PCR amplified using primers from the FOXM1 promoter region. Cyclin D1 promoter served as the positive control. Actin served as positive control for Pol II. DNA extract (10% without ChIP) was used as input. The protocol was described earlier^43,52^. The sequences of the primers are given in the Supplementary file 1.

### Tissue histology and immunohistochemical (IHC) analysis

Human patient samples (5 μm thick sections) were prepared from paraffin-embedded blocks. IHC was performed as described formerly^49,53^. Scoring was done through a semi-quantitative method whereby an overall H-score^54^ was calculated based on the degree of staining (0–100%) multiplied by the staining intensity pattern (1: negative or weak, 2: moderate, and 3: strong). The images were captured at 200X magnifications by EVOS XL Cell Imaging System (Life Technologies). Mayer’s Hematoxylin (#MHS1; Sigma, St. Louis, MO, USA) was used to counterstain the nucleus. The primary antibodies p68 (Abcam); FOXM1, Survivin (Cell Signaling Technology), and β-catenin (Santacruz Biotechnology) were used.

### Cell viability and wound healing assays

In Wound healing (scratch) assay, cells were seeded and transfected into 35 mm culture dishes until they reached 80-90% confluency after 24 h and 48h of the experiment. Fine scratches were made using sterile tips. The assay using MTT was performed as described before^50,55^. Cells were observed under 10X objective to capture all images.

### Survival assay

Colony-formation assays were conducted as described earlier^45,50^. Briefly, HCT 116 cells were cultured in 35mm plates for treatment or transfection. After 15 days, fixation and crystal violet staining was performed. The number of colonies were counted and photographed^56^ using the Nikon D5600 DSLR. The graphical and statistical analyses were done using GraphPad Prism software.

### Cell cycle analysis

HCT 116 cells were transfected or treated according to the experimental needs. After 24 or 48hrs, cells were trypsinized, ethanol (70%) fixed and processed for cell-cycle analysis. Analysis was done in BD LSR-Fortessa using FACS-Diva software (BD Biosciences)^57^.

### Statistical and bioinformatic analysis

Bioinformatics was done using UALCAN database. The student’s t-test (unpaired) was performed using GraphPad QuickCals (http://www.graphpad.com/quickcalcs/index.cfm). Manual calculation of H-scores was done. Mann–Whitney U-test was done using R module Wilcoxon-Mann-Whitney Test calculator (https://www.wessa.net/rwasp_Reddy-Moores%20Wilcoxon%20Mann-Witney%20Test.wasp). Box plots were generated using the web tool BoxPlotR (http://boxplot.tyerslab.com/). Spearman’s rank correlation coefficient (rs) was calculated using R module Spearman Rank Correlation calculator (http://www.wessa.net/rwasp_spearman.wasp/). The significance value of p<0.05 was considered to be statistically significant (*), p<0.01, p<0.001, and p<0.0001 were considered to be very significant (**), highly significant (***), and extremely significant (****), respectively.

## Acknowledgments

Authors sincerely acknowledge Dr. Kiran Kumar Naidu, Dr. Moumita Sarkar and Dr. Veenita Khare (ex-students of Dr. Mrinal K Ghosh) for their technical help in developing this project and current findings. Veenita Khare helped in the preparation of IHC tissue slides and histological experiments. We would like to thank Dr. Uttara Chatterjee (Park Clinic, Kolkata, India) for providing the human normal colon and colon carcinoma samples.

## Availability of data and materials

Data supporting the conclusion of the current study is included in the manuscript and the supplementary files.

## Ethics approval and consent to participate

All clinical and ethical regulations were followed for collecting the samples, including patient consent, and with formal approval from the ethical committee of both CSIR-IICB and Park Clinic (source). All procedures performed in this study involving animals were approved by the Ethics Committee of CSIR-IICB.

