## Supplementary file for "The DEAD-box protein p68 and β-catenin: the crucial regulators of FOXM1 gene expression in arbitrating colorectal cancer"

### Supplementary figures and legends

**Figure S1. p68 regulated FOXM1 at the transcript level. (a)** and **(b)** HCT 116 and HT29 cells were seeded respectively on 35 mm culture dishes and transfected with control sirRNA or p68-siRNA (2µg). After 48 hrs, they were harvested for total RNA extraction and subsequent cDNA preparation was done which was used in qRT-PCR analysis. The mRNA levels of p68, FOXM1, and Cyclin D1 (positive control) genes were observed. The mean (+/-) s.d is represented by error bars; the p-values were calculated using an independent, two-tailed Student's t-test, and $p < 0.0001$ is signified as ****.

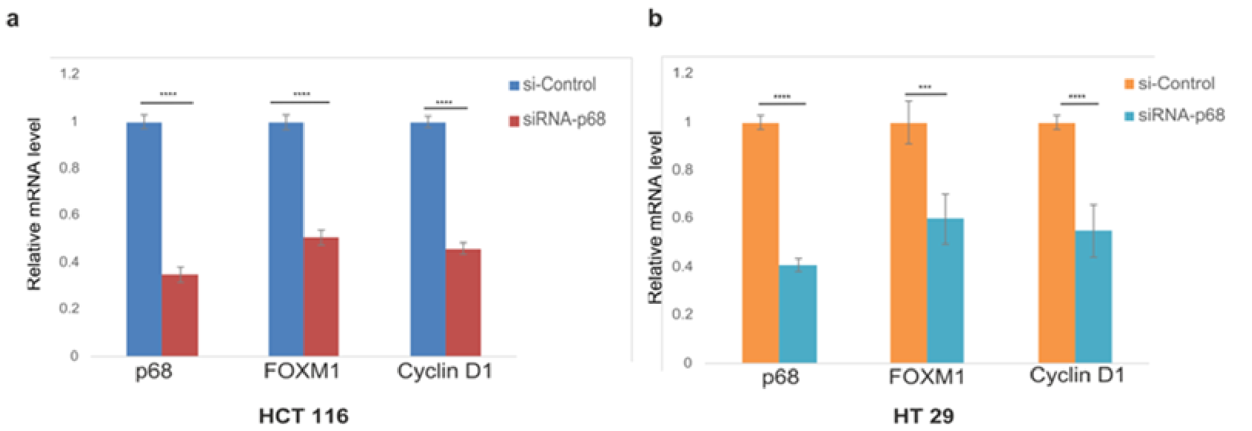


**Figure S2. FOXM1 is regulated by β-catenin.** SW480 cells were seeded on 35 mm culture dishes and transfected with empty PGZ or PGZ-β-catenin(2µg). After 48 hrs, they were harvested for total RNA extraction and subsequent cDNA preparation was done which was used in qRT-PCR analysis. The mRNA levels of β-catenin, FOXM1, and Cyclin D1 (positive control) genes were observed. The mean (+/-) s.d is represented by error bars; the p-values were calculated using an independent, two-tailed Student's t-test, and $p < 0.0001$ is signified as ****.

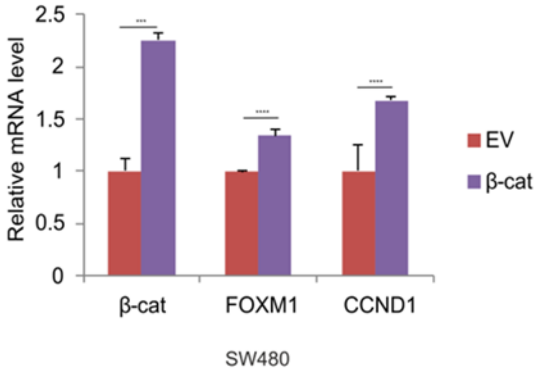

**Figure S3. Positive correlation between p68, β-catenin, FOXM1 and surviving (downstream target of FOXM1). (a)** Scatter plots representing the mean H-scores of β-catenin and Survivin in normal and colon carcinoma tissue samples, respectively. **(b)** Determination of Spearman's rank correlation coefficient ($r_s$) for positive correlation between the mean H-scores of β-catenin and Survivin in from both normal and colon carcinoma tissues. **(c)** Combined average H-scores of p68, β-catenin and Survivin were represented graphically to compare and contrast between expression profiles in normal vs colon carcinoma sample tissues. The mean (+/-) s.d is represented by error bars; the p-values were calculated using an independent, two-tailed Student's t-test, and $p < 0.0001$ is signified as ****.

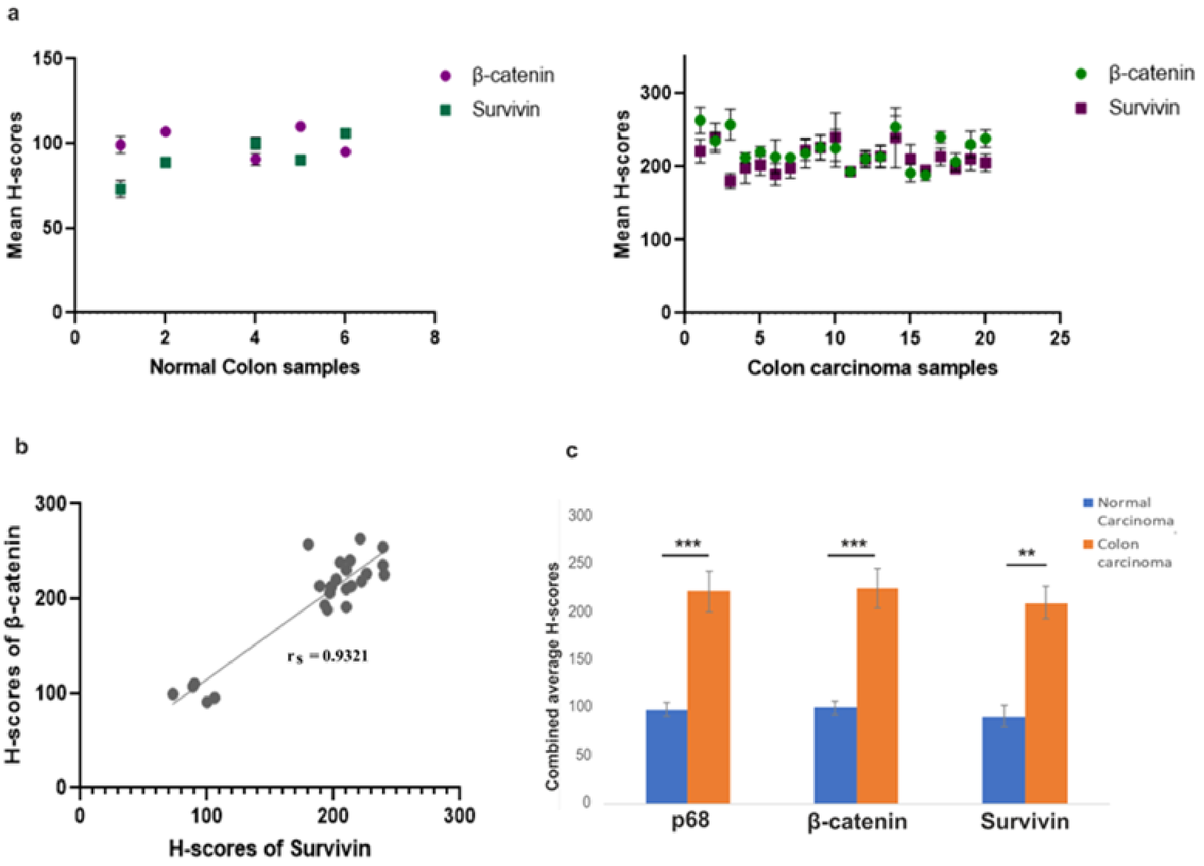

**Figure S4. p68 regulates the promoter activity of FOXM1.** Luciferase assay was conducted in HCT 116 cells wherein the cells were seeded in 35 mm culture dishes and transfected with either control siRNA or p68-siRNA along with an equivalent concentration of pGL3-WT-FOXM1-promoter construct and Renilla luciferase plasmid (50 ng for normalization). The luciferase activity was measured 48hrs post-transfection. The mean (+/-) s.d is represented by error bars; the p-values were calculated using an independent, two-tailed Student's t-test, and $p < 0.0001$ is signified as ****.

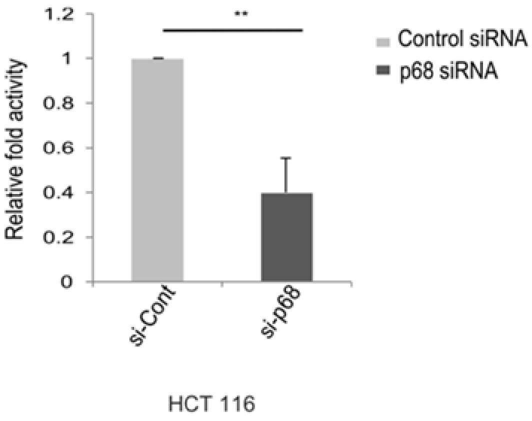

**Figure S5. Wound healing assay represent the cellular proliferation and migration by p68/FOXM1 axis.** HCT 116 cells were seeded in 35 mm cell culture dishes and were treated with Scramble or shRNA-p68 and sh-β-catenin. Later, 10ul sterile pipette tips were used to make transverse scratches on the plates and the images were captured at 0, 24, and 48 hrs. All the p-values were calculated using an independent, two-tailed Student's t-test, and $p < 0.0001$ is signified as ****.

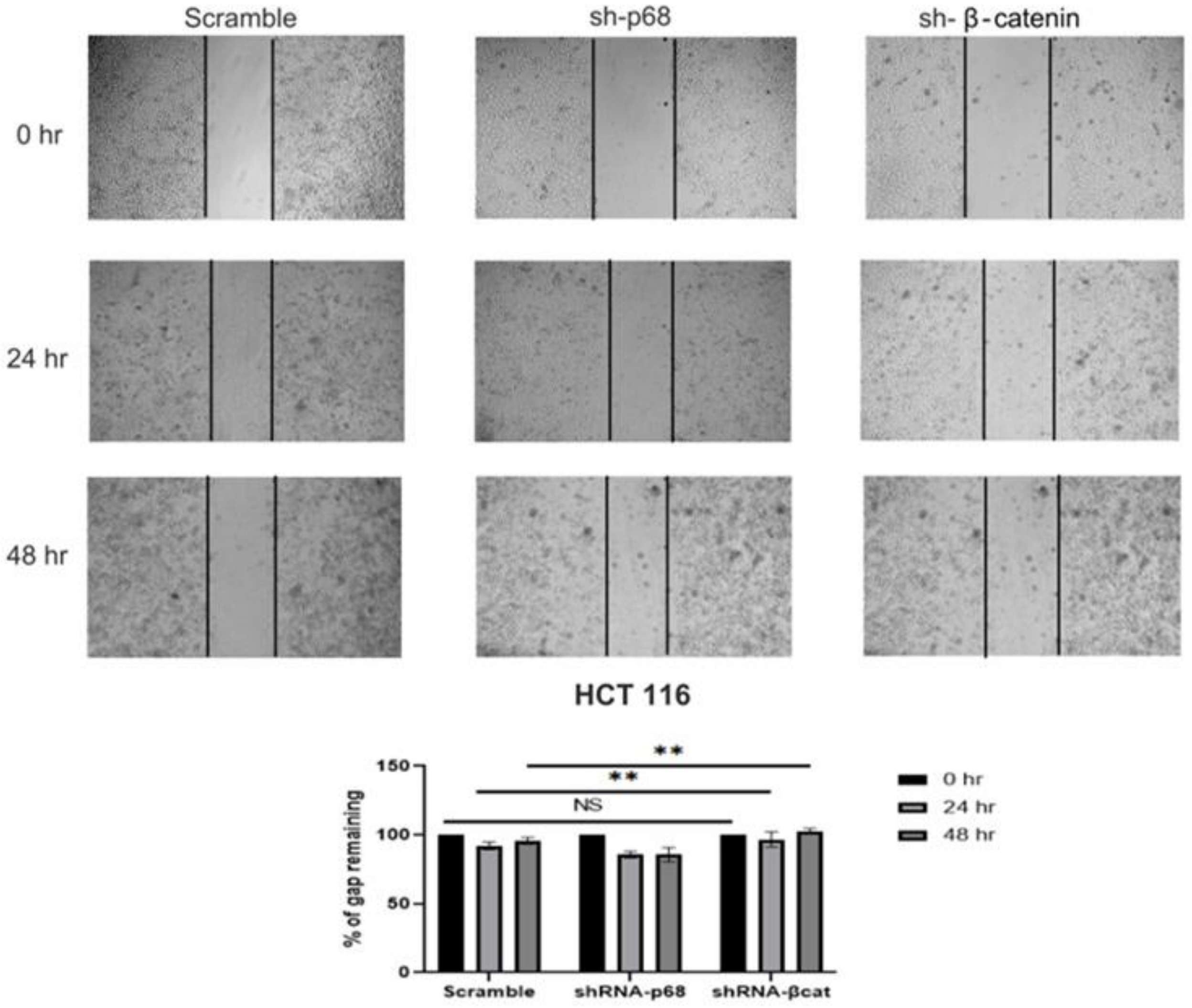

**Figure S6. p68/β-catenin synergistically regulate FOXM1 to drive oncogenesis. (a)** Cell cycle distribution profiles is a crucial functional assay of HCT 116 cells transfected with DMSO or 40μM of thiostrepton was observed and represented graphically where the bar graph signifies the change in the percentage of S phase cells in the treated compared to the control. **(b)** and **(c)** Colony formation assay was done in HCT 116 cells by fixing the cells in crystal violet stain after treatment with LiCl and also through transfection by Scramble or sh-β-catenin. The relative number of colonies was observed, counted and represented graphically. All the p-values were calculated using an independent, two-tailed Student's t-test, and $p < 0.0001$ is signified as ****.

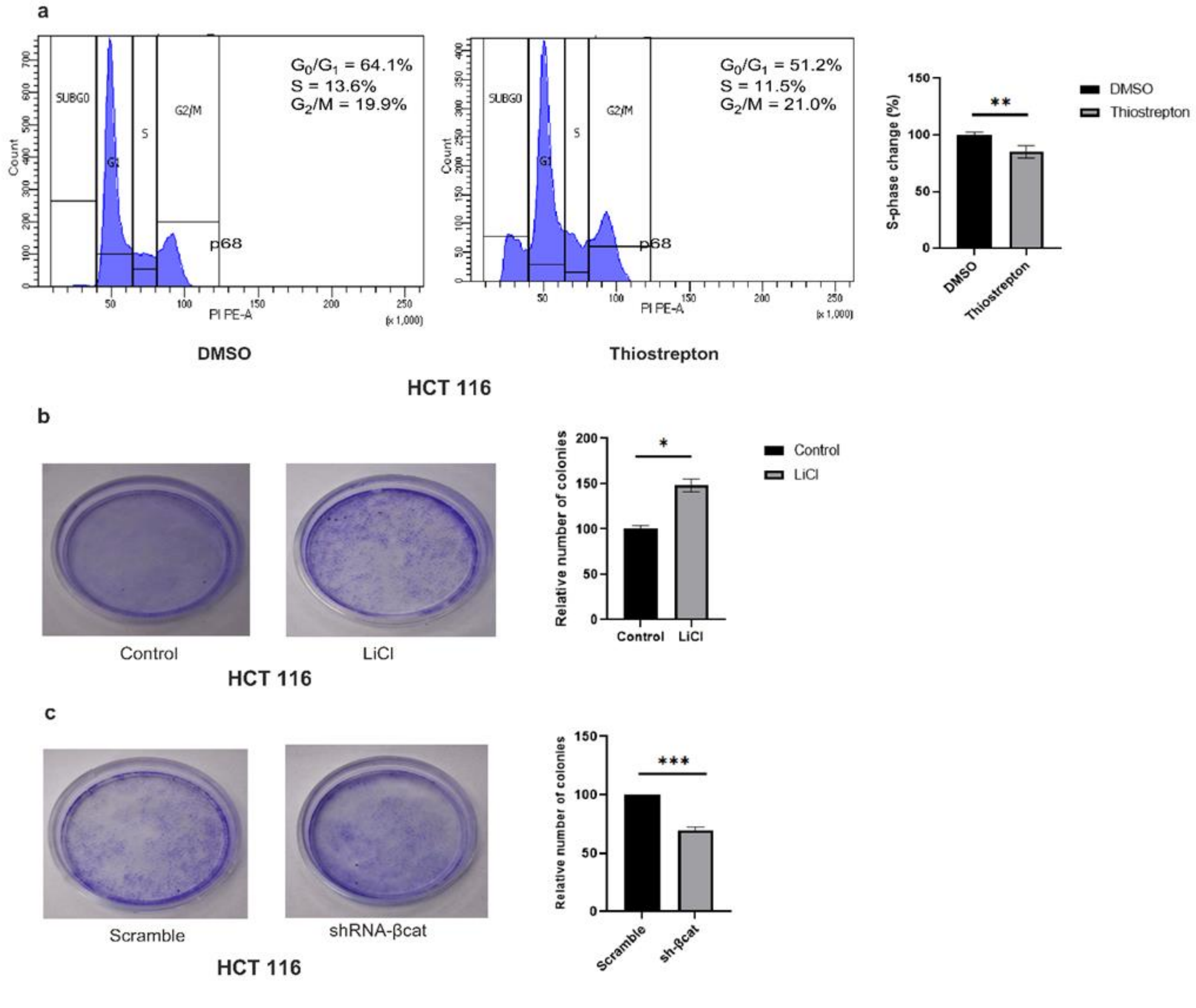
